# LncRNA-UCA1 inhibits the astrocyte activation in the temporal lobe epilepsy via regulating JAK/STAT signaling pathway

**DOI:** 10.1101/765982

**Authors:** MM Hongxin Wang, BM Guangyan Yao, MM Lei Li, MM Zhaoyin Ma, MM Jing Chen, DM Wen Chen

## Abstract

This article aimed to reveal the mechanism of Urothelial cancer associated 1 (UCA1) regulated astrocyte activation in temporal lobe epilepsy (TLE) rats via JAK/STAT signaling pathway. A model of TLE was established based on rats via kainic acid (KA) injection. All rats were divided into sham group, KA group, normal control (NC) + KA group and UCA1 + KA group. The Morris water maze was used to test the learning and memory ability of rats, and the expression of UCA1 in hippocampus was determined by qRT-PCR. Surviving neurons were counted by Nissl staining, and expression of glial cells glial fibrillary acidic protein, p-JAK1, and p-STAT and glutamate/aspartate transporter (GLAST) was analyzed by immunofluorescence and Western blot. A rat model of TLE was established by intraperitoneal injection of KA. QRT-PCR and fluorescence study showed that UCA1 inhibited astrocyte activation in hippocampus of epileptic rats. Meanwhile, the MWM analysis indicated that UCA1 improved the learning and memory in epilepsy rats. Moreover, the Nissl staining showed that UCA1 might has protective effect on neuronal injury induced by KA injection. Furthermore, the immunofluorescence and Western blot analysis revealed that the overexpression of UCA1 inhibited KA-induced abnormal elevation of GLAST, astrocyte activation of JAK/STAT signaling pathway, as well as hippocampus of epilepsy rats. UCA1 inhibited hippocampal astrocyte activation and GLAST expression in TLE rats via regulating JAK/STAT signaling, and improved the adverse reactions caused by epilepsy.

## Introduction

Epilepsy is the most common serious neurological disorder worldwide ^[1, 2]^. As a classic type of epilepsy, temporal lobe epilepsy (TLE) is a common intractable in adults characterized by recurrent and unprovoked focal seizures originating from the temporal lobe ^[3]^. Worldwide, it results in 125,000 deaths in one year ^[4]^. The surgical treatment such as resective temporal lobe surgery is the mainly strategy for patients with TPL ^[5]^. In addition, the outcomes of TLE patients underwent surgery are greatly limited by the occurrence of diverse postoperative neurological complications ^[6]^. Therefore, novel therapeutic strategies with high efficiency and low adverse effects for TLE are urgently needed.

Understanding the mechanisms of LTE at the molecular and structural level is valuable for clinical treatment ^[7]^. The long non-coding RNA (lncRNA) urothelial carcinoma-associated 1 (UCA1) is proved to participate in several diseases ^[8, 9]^. A previous study shows that there is a dynamic regulation effect of UCA1 on epilepsy rats ^[10]^. Actually, the alternation of glial cells is strongly associated with the development of LTE ^[11]^. A previous study shows that lncRNAs such as H19 promotes activation of hippocampal glial cells and serves as a therapeutic tool to prevent epileptogenesis by regulating JAK/STAT pathway ^[12]^. The formation of glial fibrillary acidic protein (GFAP)-positive glia is realized by activation of JAK/STAT3 pathway ^[13]^. Although sporadic researches prove the relations among UCA1, JAK/STAT3 pathway and epilepsy, whether UCA1 take part in the LTE progression via JAK/STAT signaling pathway is still not fully revealed. Glutamate aspartate transporter (GLAST) is expressed throughout the central nervous system, and is highly expressed in astrocytes and Bergmann glia in the cerebellum ^[14]^. Previous studies show that the up-regulation of GLAST is closed related with the development of epilepsy ^[14, 15]^. However, the specific roles of UCA1 on GLAST during the pathological changes of TLE is still unclear.

In this study, a model of TLE was established based on male Sprague-Dawley (SD) rats. Based on the TLE model, the Morris water maze (MWM) analysis, qRT-PCR analysis, Nissl staining, Immunofluorescence analysis, as well as Western blot analysis were investigated. Our findings may reveal the detail roles of UCA1 on neural damages of TLE, and reveal novel mechanisms responsible for the treatment of TLE.

## Material and methods

### TLE rats model construction

Male SD rats (200-220 g, 6-8 weeks, SPF grade) were purchased from Beijing Weitonglihua Laboratory Animal Technology Co., Ltd. Rats were fed in a standalone environment at 22°C and 50% relative humidity under the alternating day and night of 12 h/12 h with free access to water and food. Then, the AAV9 vector carrying UCA1 (AAV9-UCA1, Shanghai Jikai Gene Chemical Technology Co., L) and an AAV empty vector (AAV-NC) were injected into right dorsal and ventral hippocampus of rats. A total of 2nmol kainic acid (KA, Sigma-Aldrich, MO, USA) was injected into right dorsal hippocampus of rats (KA group, NC + KA group and UCA1 + KA group) at 14 day of AVV vector injection until reaching IV grade epilepsy as proposed by Racine ^[16]^. Rats injected with an equal amount of physiological saline instead of the KA were considered to be the sham group. Subsequently, on the 10th day of successful establishment of epilepsy model, the brain of rats were obtained after anesthetized with 10% chloral hydrate (5 mL/kg), followed by the cerebellum and olfactory bulb tissue harvest. Finally, the dorsal hippocampus was coronally sectioned (25 μm) for further investigation. This study was approved by the local ethics committee, and all experiments were in accordance with the guide for the care and use of laboratory animals established by United States National Institutes of Health (Bethesda, MD, USA).

### MWM analysis

After induction of status epilepticus, rats in each group were trained in the Morris water maze for 7 days to assess their spatial learning and memory ability. Briefly, four points along the perimeter of the maze arbitrarily designated as starting points where the mice were released, facing the wall of the tank, at the beginning of each trial. Statistical parameters of mice in water maze was recorded. Other task parameters remained identical to the acquisition procedures.

### QRT-PCR assay

Total RNA was extracted from the hippocampus of rats using SYBR Premix Ex Taq kit (Takara Biotechnology Co. Ltd., Dalian, China), and reverse-transcribed by FastQuant RT Kit (Tiangen, Beijing, China) in accordance with manufacturers’ instructions. The qRT-PCR was performed on ABI PRISMR 7300 Sequence Detection System (Applied Biosystems, Foster City, CA, USA). In detection of UCA1 expression (UCA1 forward, 5’-ACCTCAACCCAAAGGCAGTC-3’; UCA1 reverse, 5’-GCCTTTGTGCCGCTACTTTT-3’), the PCR program included 95°C for 10 s, 40 cycles of 95°C for 5 s, 60°C for 15 s and 72°C for 31 s. GAPDH was used as an internal reference (GAPDH forward: 5’-CGGAGTTGTTCGTATTCGG-3’; GAPDH reverse: 5’-TACTAGCCGATGATGGCATT-3’)., and the data was analyzed by the 2^−ΔΔCt^ method ^[17]^, and all oligonucleotide primers were designed and synthesized by Takara (Takara Biotechnology Co. Ltd., Dalian, China).

### Nissl staining

The dorsal hippocampal frozen coronal sections were stained with 1% tolridine blue (Shanghai Shenggong Bioengineering Co., Ltd.), gradient dehydration of alcohol, transparent by xylene and then sealed with a neutral gum. A Nikon Eclipse 80i microscope (Nikon, Japan) was used for photographing, and Leica Qwin Analysis software V2.8 (Leica Microsystems, Germany) was used to count the number of viable cells.

### Immunofluorescence analysis

Paraffin sections of hippocampus were dewaxed and rehydrated with ethanol, followed by antigen retrieval in 3% hydrogen peroxide for 10 min. Then sampels were treated with 5% goat serum occlusion and primary antibodies (neuron nuclei (NeuN), ab177487, 1:500, Abcam, Cambridge, rabbit anti-rat GFAP, ab7260, 1:500, Abcam, Cambridge, MA, USA; rabbit monoclonal anti-p-Stat3, ab76315, 1:500, Abcam, Cambridge, MA, USA; rabbit polyclonal anti-EAAT1 (GLAST), #5684, 1:100, Cell Signaling Technology, Beverly, MA, USA). Then, the samples were treated with secondary antibody Alexa Fluor 488-labeled goat antibody (Merck Biosciences, Nottingham, UK). Finally, the samples were stained for 10 minutes by the addition of DAPI (Sigma, Missouri, USA), followed by images capture and cells counting.

### Western blot

Total proteins extracting from hippocampus tissue were separated by electrophoresis on 10% polyacrylamide gels and placed on polyvinylidenefluoride (PVDF) membranes. After blocked with 5% skim milk - TBST solution for 1 h, the primary antibodies including rabbit polyclonal anti-GFAP (ab7260, 1:1000, Abcam, Cambridge,MA, USA), rabbit monoclonal anti-p-JAK1 (#74129, 1:1000, Cell Signaling Technology, Beverly, MA, USA), rabbit monoclonal anti-JAK1 (#3344, 1:1000, Cell Signaling Technology, Beverly, MA, USA), rabbit monoclonal anti-p-Stat3 (ab76315, 1:1000), rabbit monoclonal anti-Stat3 (ab68153, 1:1000) and rabbit polyclonal anti-EAAT1 (ab416, 1:1000, Abcam, Cambridge, MA, USA) were performed on membrane. Rabbit monoclonal anti-GAPDH (Abcam, ab181602, 1:3000) was used as internal reference. Then the membrane was treated with HRP-labeled corresponding secondary antibody. Finally, protein brands were visualized using Gel imaging system (Thermo Fisher Scientific).

### Statistical analysis

All data were expressed as mean ± standard deviation (SD). Statistical analysis was performed by SPSS 21.0 (SPSS, Inc., Chicago, IL, USA). The differences between groups was revealed using a Student’s t-test. One-way ANOVA for repeated measures followed by Tukey’s multiple comparisons test. The P < 0.05 was selected as threshold for significantly different.

## Results

### LncRNA UCA1 inhibited epilepsy in rats with epilepsy

Compared with sham group, the seizure frequency of rats in KA group increased significantly (P < 0.001), and the frequency of seizures of rats in the UCA1+KA group was lower than that of NC+KA group (P < 0.05) (Figure 1A). The UCA1 expression in KA-induced epileptic rats was significantly decreased (P < 0.01, Figure 1B). Compared with AVV-NC group, the UCA1 expression was significantly higher in AVV-UCA1 group (P < 0.01, Figure 1C). Fluorescence observation of brain sections revealed that the injected vector was fused to the hippocampus (Figure 1D).

**Figure 1.**
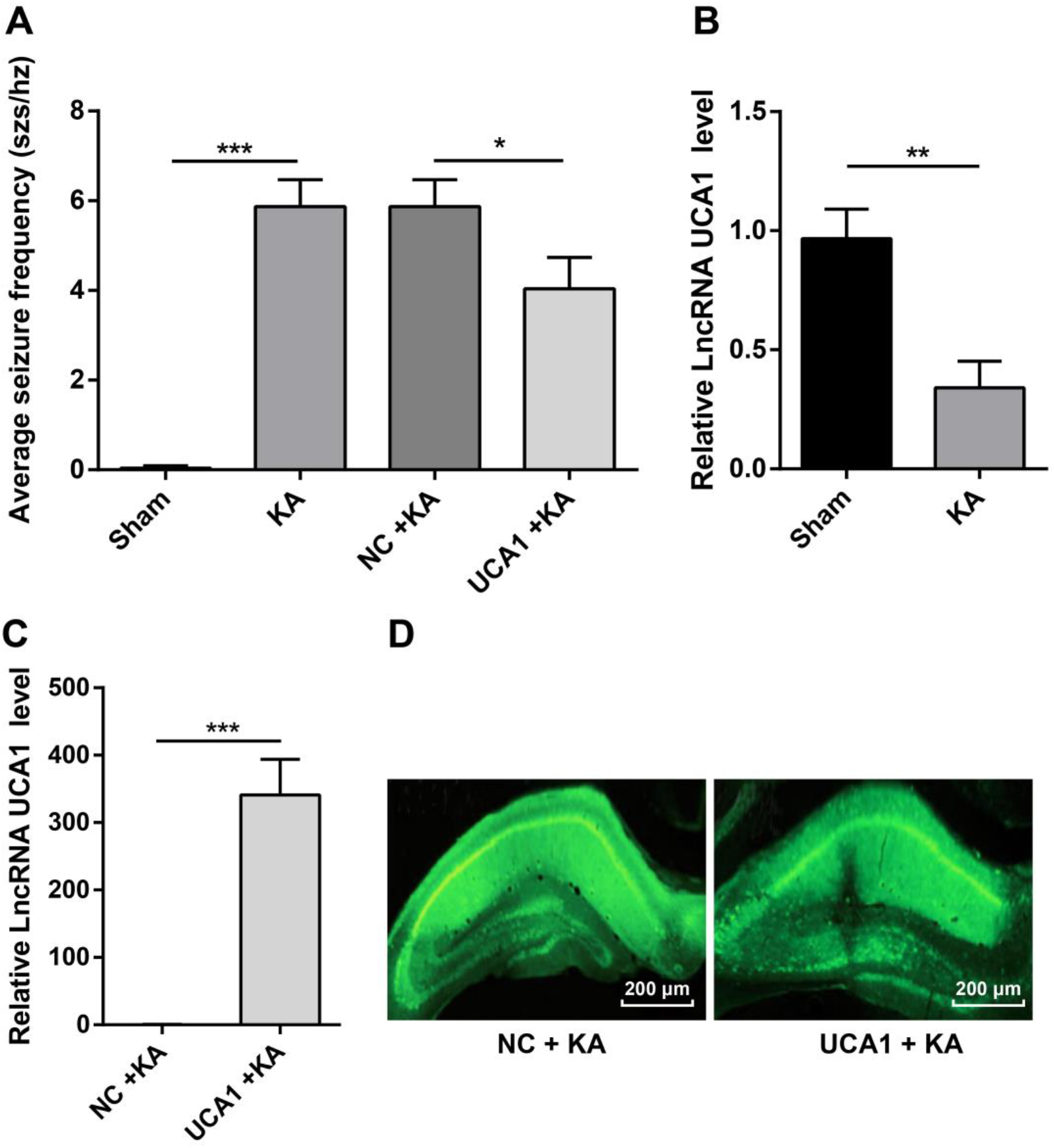
Levels of lncRNA UCA1 in each group of rats. A, the frequency of seizures in each group of rats on the 10th day; B, lncRNA UCA1 levels in normal and epileptic rats were detected by qRT-PCR; C, the level of lncRNA UCA1 in epileptic rats injected with AAV vector was detected by qRT-PCR; D, the results of immunofluorescence in hippocampus of epileptic rats injected with AAV vector (200μm). *P < 0.05, **P < 0.01 and ***P < 0.001, versus the sham group.

### LncRNA UCA1 improved learning and memory in epilepsy rats

The results obtained in MWM assay showed that the latency of KA group was significantly longer than that of Sham group, and that of UCA1 + KA group was significantly shorter than that of NC + KA group (Fig. 2A). (both P <0.001, Figure 2A). However, there was no significant difference in the speed of swimming between the groups (P > 0.05, Figure 2B). Compared with Sham group, the swimming distance of rats in KA group increased significantly, while that in UCA1 + KA group was shorter than that in NC + KA group (Figure 2C). Simultaneously, overexpression of UCA1 in epileptic rats took more time in the target area and traversed the platform than in NC + KA group. It was suggested that lncRNA UCA1 had the function of improving learning and memory in epilepsy rats (Figure 2D, 2E).

**Figure 2.**
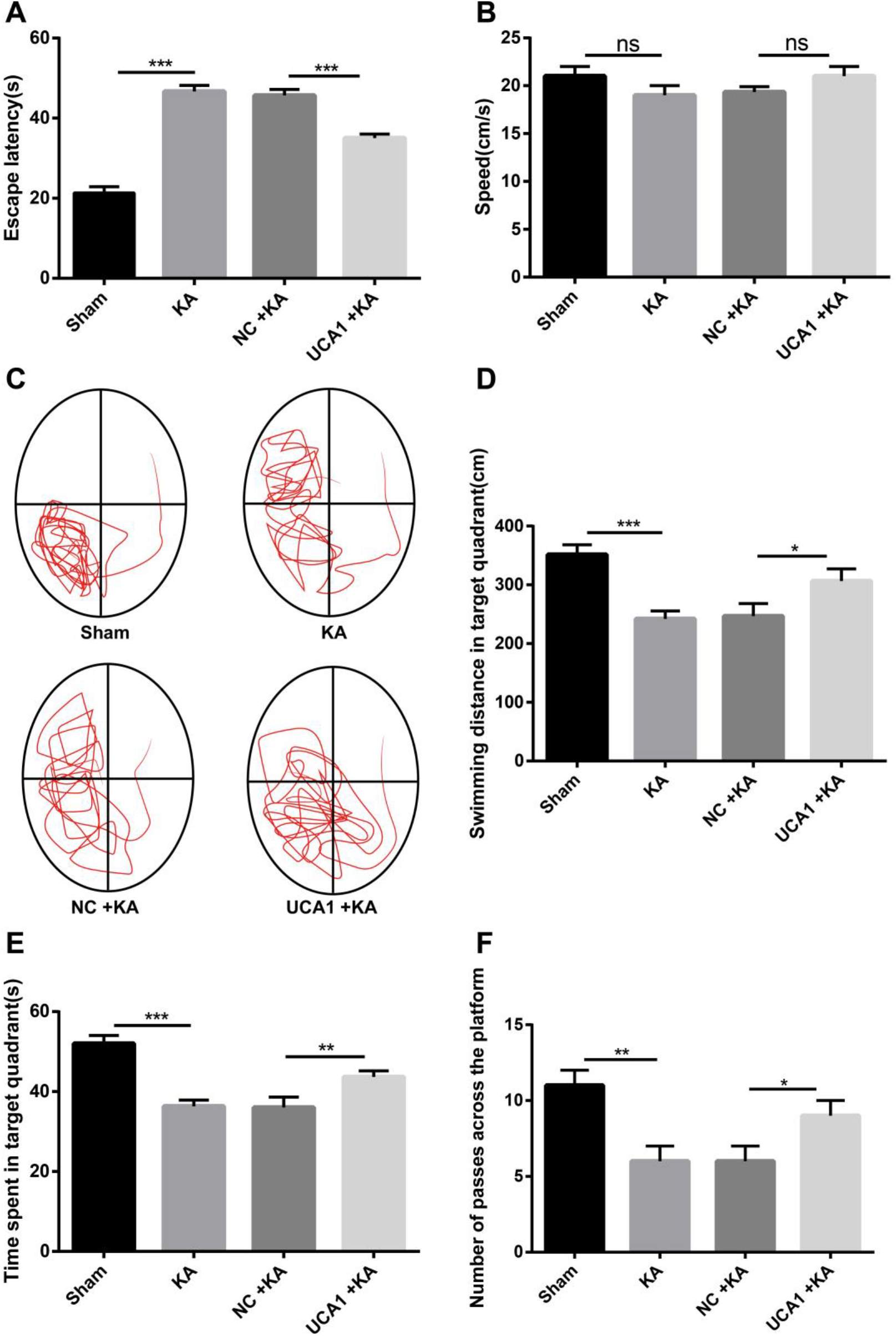
lncRNA UCA1 improved learning and memory in epilepsy rats. A, the incubation period of rats in each group in the positioning navigation test on the fifth day of training; B, swimming speed of each group of rats in the space exploration test; C, path of the rat when it finds the target quadrant of the platform; D, the residence time of rats in the quadrant of the platform; E, number of times the rats in each group crossed the platform. *P < 0.05, **P < 0.01 and ***P < 0.001, versus the sham group.

### LncRNA UCA1 attenuated hippocampal neuronal injury in epilepsy rats

Results of the hippocampal coronal section stained by Nissl were shown in Figure 3. In the sham group, the pyramidal cells in the CA3, CAl and hilar areas were blue, dark-stained, and abundant with the Nissl bodies in the cytoplasm. In KA group and UCA1 + KA group, neurons in CA3, CAl area and portal area were disordered and incomplete, showing dense stained cell fragments, a small number of surviving cells, cell body shrinkage, nuclear pyknosis and cytoplasmic Nissl body reduction. In UCA1 + KA group, there were more neurons in CA3, CAl and portal areas, and the layers were dense but not disordered. These results suggest that LncRNA UCA1 has protective effect on neuronal injury induced by KA injection.

**Figure 3.**
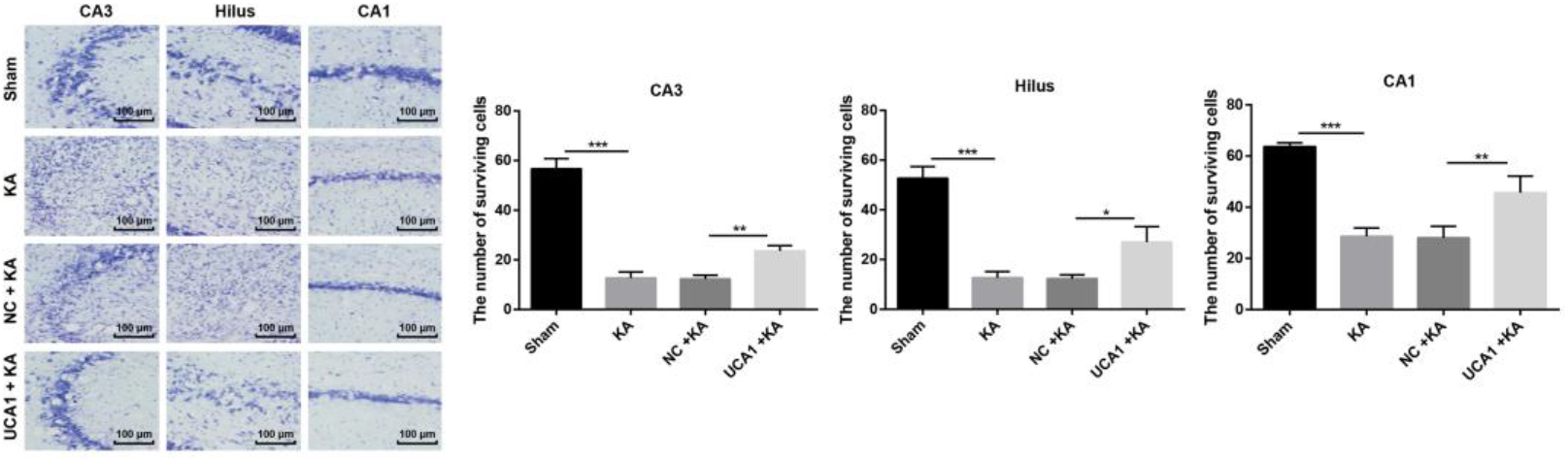
Effect of lncRNA UCA1 on hippocampal neuronal injury of epileptic rats. The arrow on the left indicated CA3, CAl and portal areas (scale = 100 um). The enlarged image corresponded to the left-most marked box (scale = 100 um). On the right was the number of surviving neurons in CA3, CAl and portal areas. *P < 0.05, **P < 0.01 and ***P < 0.001, versus the sham group.

### The effect of UCA1 overexpression on epilepsy rats

The results of immunofluorescence staining of GFAPd were shown in Figure 4A. Compared with the sham group, GFAP in the hippocampal CA3 region of epileptic mice was highly expressed. However, results of UCA1 + KA group showed that overexpression of UCA1 inhibited astrocyte activation in hippocampus of epileptic rats, which was confirmed by GFAP quantitative analysis and Western blot (Figure 4B, 4C).

**Figure 4.**
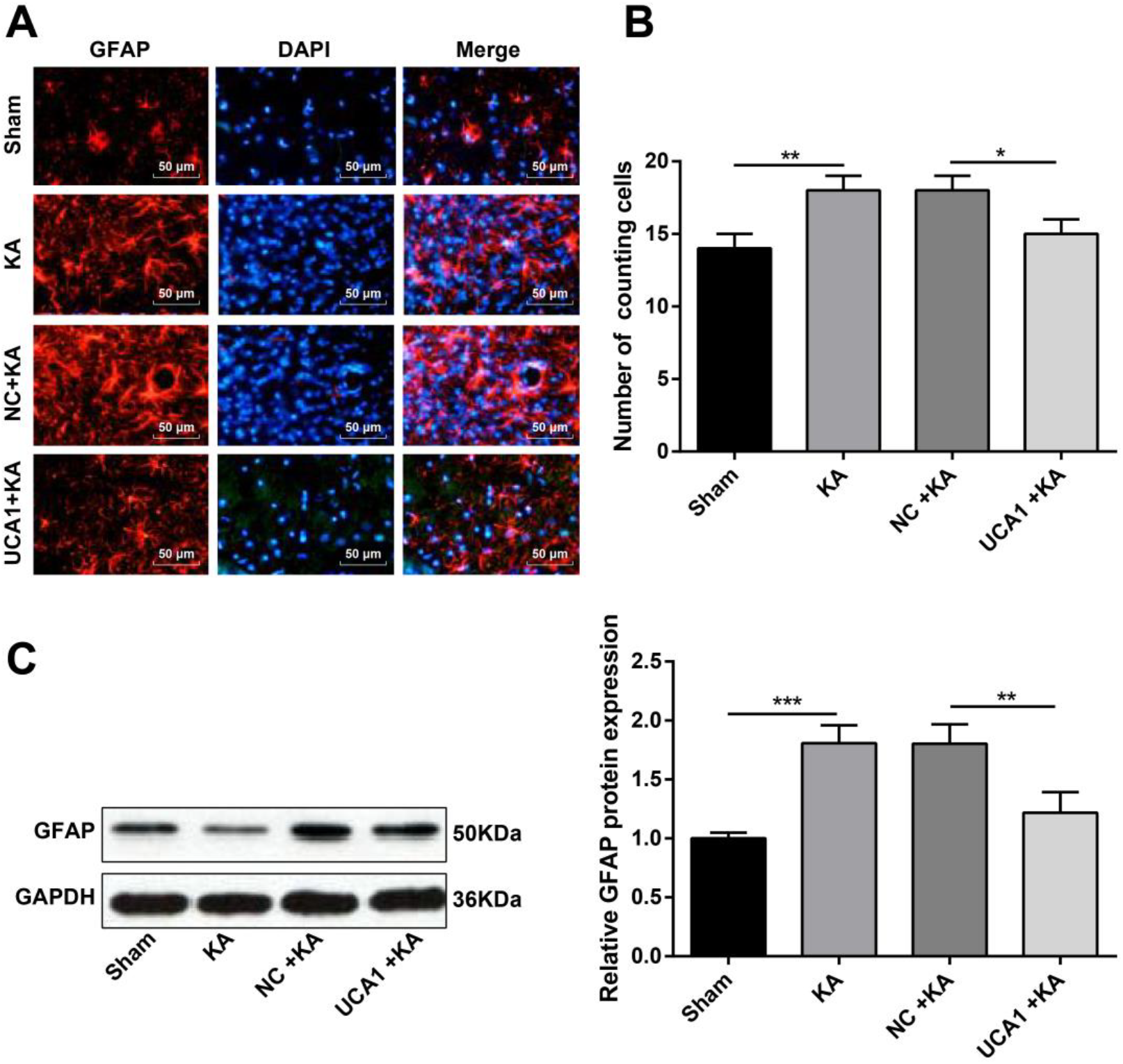
Overexpression of LncRNAUCA1 inhibited astrocyte activation in the hippocampus of epileptic rats. A, photomicrographs of GFAP expression in the hippocampus of rats in each group (Scale = 50 μm); B, number of GFAP fluorescent cells in the hippocampal CA3 region of the KA injection side; C, levels of GFAP protein in the hippocampal CA3 region of each group. *P < 0.05, **P < 0.01 and ***P < 0.001, versus the sham group.

### Overexpression of lncRNA UCA1 inhibited JAK/STAT signaling pathway

The cytoplasm of GFAP-positive cells was green star-shaped, and the nucleus of p-STAT3-positive cells was red round or oval, both of which were mainly distributed in the hippocampal CA3 area and the portal area. The coincidence of GFAP-positive green cells and p-STAT3-positive red cells indicates that STAT3 activation occurs in astrocytes during epilepsy (Figure 5A). More importantly, the p-JAK1 and p-STAT3 expression in the KA group were promoted compared with the sham group (all P < 0.01), whereas up-regulation of UCA1 inhibited the expression of p-JAK1 and p-STAT3 (Figure 5B, 5C). This result indicated that overexpression of UCA1 inhibited KA-induced activation of JAK/STAT signaling pathway.

**Figure 5.**
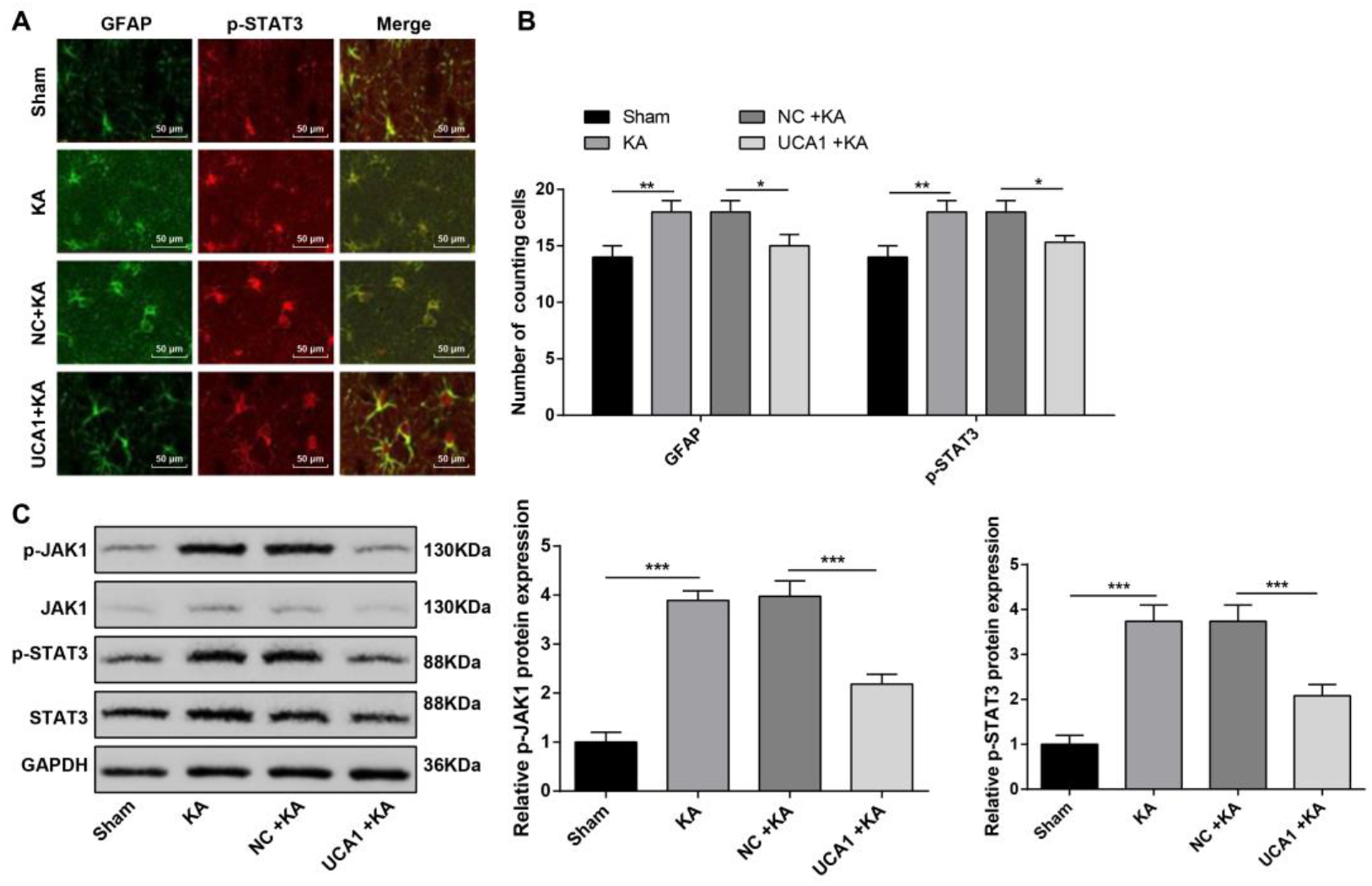
Overexpression of LncRNAUCA1 inhibited JAK/STAT signaling pathway. A, the activation of STAT3 in astrocytes detected by immunofluorescence (scale = 50 μm); B, number of GFAP and p-STAT3 fluorescent cells in the hippocampal CA3 region of the KA injection side; C, the levels of p-JAK1 and p-STAT3 proteins in each group were detected by Western blot. *P < 0.05, **P < 0.01 and ***P < 0.001, versus the sham group.

### Overexpression of lncRNA UCA1 inhibited GLAST expression in hippocampal astrocytes

GLAST and GFAP were co-localized by immunofluorescence double staining and photographed under a microscope as shown in Figure 6A. Immunofluorescence and Western blot showed that compared with the Sham group, the expression of GLAST in KA group was increased dramatically (P < 0.01), and UCA1 overexpression inhibited GLAST expression (Figure 6 B-C). These results suggested that UCA1 overexpression inhibits KA-induced abnormal elevation of GLAST.

**Figure 6.**
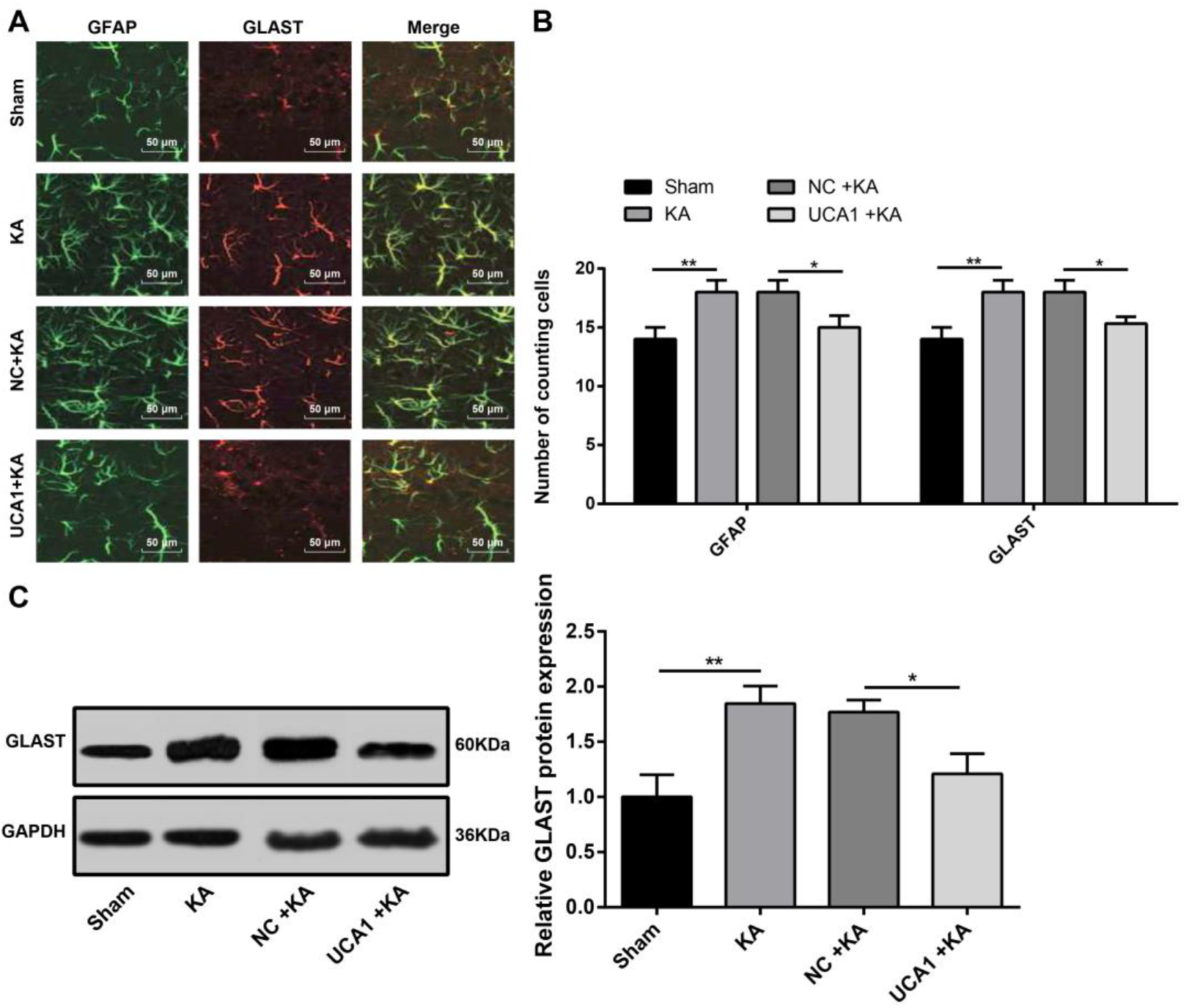
Overexpression of LncRNAUCA1 inhibited GLAST expression in hippocampal astrocytes. A, GLAST localized to astrocytes (scale bar = 50 μm); B, number of GFAP and GLAST fluorescent cells in the hippocampal CA3 region of the KA injection side; C, the levels of GLAST proteins in each group detected by Western blot. *P < 0.05, **P < 0.01 and ***P < 0.001, versus the sham group.

## Discussion

Further understanding of molecular mechanism during TLE progression is benefit for novel treatment strategy investigation. In current study, a rat model of TLE was established by intraperitoneal injection of KA. The qRT-PCR and fluorescence study showed that UCA1 inhibited astrocyte activation in hippocampus of epileptic rats. Meanwhile, the MWM analysis indicated that UCA1 improved the learning and memory in epilepsy rats. Moreover, the Nissl staining showed that UCA1 might has protective effect on neuronal injury induced by KA injection. Furthermore, the immunofluorescence and Western blot analysis revealed that the overexpression of UCA1 inhibited KA-induced abnormal elevation of GLAST, astrocyte activation of JAK/STAT signaling pathway, as well as hippocampus of epilepsy rats.

UCA1 play a key role in the development and progression of a variety of human tumors ^[18]^. The expression of lncRNA UCA1 was a dynamic change in the process of epilepsy, which was involved in the pathogenesis of epilepsy ^[10]^. Actually, the astrocytes are considered to be a key factor in the induction of epilepsy ^[19]^. Previous study shows that astrocytic dysfunction was the cause of seizure activity or the transmission ^[20]^. Astrocytes plays a crucial role in the development of epilepsy, and astrocyte dysfunction occurs early in the epileptic state ^[21]^. A previous study shows that lncRNA play important roles in the activation of rats astrocytes ^[22]^. Despite of UCA1, GLAST is proved to be dysregulation in the radial glia-astrocyte lineage of developing mouse spinal cord ^[23]^. Glutamate is the major excitatory neurotransmitter in the central nervous system and vital for most aspects of normal brain function ^[24]^. Genetic manipulations associated with the expression of GLAST can induce epileptic syndrome or increase seizure thresholds ^[25]^. In the current study, GLAST in the hippocampus of rats was dramatically increased than that in normal rats, but inhibited by UCA1. In addition, the qRT-PCR and fluorescence study showed that UCA1 inhibited astrocyte activation in hippocampus of epileptic rats. Thus, we speculated that the overexpression of UCA1 might take part in the TLE progression via inhibiting astrocyte activation and GLAST expression in hippocampus of epileptic rats.

Epileptic induction of GFAP expression is the first step in the response to epileptic induction of astrocyte hypertrophy ^[26]^. Hu et al. shows that the expression of GAFP is increased after chronic epileptic seizures ^[27]^. Samuelsson et al. indicates that the level of GFAP increased with time in epilepsy model, and the level of extracellular glutamate increase during epilepsy due to increase synaptic release and disturbance of glutamate uptake ^[28]^. Actually, the activation of STAT3 and up-regulation of GFAP expression are involved in the process of epilepsy ^[29]^. A previous study shows that the formation of GFAP-positive glia is based on the activation of the JAK/STAT3 pathway ^[13]^. Inhibition of the JAK/STAT pathway reduced the expression of glutamate transporters in astrocytes and down-regulated the expression of GLAST ^[30]^. Interestingly, lncRNA UCA1 play an vital role in the biological function of pathway involved by STAT ^[31]^. In this study, the expression of GFAP and p-STAT3 in the CA3 region of the hippocampus of KA-induced epilepsy was significantly higher than that of non-epileptic rats, and the expression of GFAP and p-STAT3 in the hippocampus of rats with epilepsy treated with lncRNA UCA1 was significantly decreased. Thus, we speculated that the overexpression of UCA1 might inhibit KA-induced activation of JAK/STAT signaling pathway and decreased expression of GFAP. However, to understand the mechanism of UCA1 on the function of epilepsy requires in-depth knowledge on the targets of UCA1 and a larger scale of UCA1-target gene screening. In this study, this knowledge was fragmented and still needed a further investigation.

## Conclusions

In conclusion, lncRNA UCA1 inhibited hippocampal astrocyte activation and GLAST expression in TLE rats via regulating JAK/STAT signaling, and improved the adverse reactions caused by epilepsy.

## Supplementary Materials

None.

## Acknowledgments

None.

## Authors’ contributions

Hongxin Wang, Guangyan Yao and Lei Li designed and analyzed the experiment, and were major contributors in writing the manuscript. Zhaoyin Ma, Jing Chen and Wen Chen performed the experiment. All authors read and approved the final manuscript.

## Conflict of interest

The authors declare that they have no conflicts of interest with the contents of this article.

